# FSP1 stem/progenitor cells are essential for TMJ growth and homeostasis

**DOI:** 10.1101/2024.08.16.608236

**Authors:** Ticha Tuwatnawanit, Wilma Wessman, Denisa Belisova, Zuzana Sumbalova Koledova, Abigail S Tucker, Neal Anthwal

## Abstract

The temporomandibular joint (TMJ) is one of the most used joints in the body. Defects and wear in the cartilage of the joint, condyle, and fibrocartilage disc lie at the heart of many common TMJ disorders. During postnatal development the condyle acts as a growth centre for the mandible with cells moving as a conveyor belt away from the top of the condyle as they differentiate. The superficial layers of the condyle have been proposed to contain stem/progenitor populations to allow growth and maintain homeostasis. Here we have focused on the role of FSP1 (also known as S100a4), as a key fibroblast stem/progenitor marker for the condyle. Lineage tracing using *FSP1-Cre;mTmG* mice revealed that FSP1-positive cells were restricted to the superficial fibroblast zone giving rise to all layers of the condyle over time. The FSP1-expressing cells overlapped with other putative stem cell markers of the condyle, Gli1 and Scleraxis. BrdU pulse chase experiments highlighted that a subset of FSP1 fibrocartilage were label-retaining, suggesting that FSP1 labels a novel stem/progenitor cell population in the condyle. Destruction of FSP1-expressing cells by conditional diphtheria toxin activity in *FSP1-Cre;DTA* mice resulted in severe TMJ osteoarthritis (TMJOA) with loss of the cartilage structure. Lgr5 expressing cells in the superficial layer of the condyle, have previously been shown to create a Wnt inhibitory niche. FSP1 expression postnatally was associated with a reduction in canonical Wnt activity in the condyle. Importantly, constitutive activation of Wnt/β catenin in FSP1 cells, led to a downregulation of FSP1 and progressive postnatal loss of TMJ condylar hyaline cartilage due to loss of the superficial stem/progenitor cells. These data demonstrate a novel role for FSP1-expressing cells in the superficial zone in growth and maintenance of the TMJ condylar cartilage and highlight the importance of regulating Wnt activity in this population.

## Introduction

The temporomandibular joint (TMJ) is one of the most frequently utilized joints in the human body, comprising the condylar process of the mandible (lower jawbone) and the glenoid fossa of the temporal bone (upper jaw). A fibrous disc within a synovial capsule lies between these structures, serving as a cushioning element. TMJ articular surface is covered with fibrocartilage, a dense and avascular connective tissue rich in collagen types I and II, which resembles a mix of fibrous and hyaline cartilage (Benjamin and Ralphs, 2004; Delatte et al., 2004; Lowe and Almarza, 2017). This fibrocartilage provides a functional buffer between the bony surfaces of the condyle and fossa, accommodating the extensive range of TMJ movements (Wadhwa and Kapila, 2008).

Temporomandibular joint osteoarthritis (TMJOA) poses a significant clinical challenge due to the erosion of TMJ cartilage. The limited regenerative capacity of fibrocartilage leads to degradation of the condyle and fossa articular cartilage, inflammatory subchondral bone remodelling, synovitis, extracellular matrix (ECM) degradation, and clinical symptoms including pain, joint noises, and impaired jaw function (Cardoneanu et al., 2022; Wang et al., 2015). TMJOA, the most severe form of temporomandibular disorder (TMD), predominantly affects women and accounts for over half of TMD cases (Alzahrani et al., 2020).

The condylar process develops as a secondary cartilage during embryogenesis, integrating with the dentary bone and serving as a growth centre in postnatal life (Frommer, 1964). The mature mandibular condyle is structured into four zones: a superficial fibrous tissue, a prechondroblastic zone (expressing SOX9), a chondroblastic zone (expressing Collagen II), and a hypertrophic zone (expressing Collagen X) (Shibukawa et al., 2007), with cells moving as a conveyer belt through the zones as they differentiate. The superficial fibroblast tissue, which forms the articular surface and interfaces with the disc, has been suggested to house a stem cell population with canonical Wnt signalling crucial for maintaining fibrocartilage homeostasis (Embree et al., 2016; Ruscitto et al., 2023). Lineage tracing has highlighted a number of putative stem cell markers, including *Gli1* and *Scleraxis* (Ma et al., 2021; Lei et al., 2022)

Fibrocartilage, lining the TMJ articular surfaces, has unique molecular features compared to hyaline or elastic cartilage, notably high levels of collagen I and the expression of fibroblast-specific protein 1 (FSP1), also known as S100A4 (Delatte et al., 2004; Park et al., 2015). FSP1, a member of the S100 family of calcium-binding proteins, plays a crucial role in various diseases, including fibrosis, cirrhosis, pulmonary diseases, cardiac hypertrophy, neuronal injuries, and cancer (Strutz et al., 1995; Schneider, Hansen, and Sheikh, 2008). These conditions commonly involve fibrosis, tissue remodelling, epithelial-mesenchymal transition, and inflammatory mechanisms (Schneider, Hansen, and Sheikh, 2008). FSP1 is expressed in a variety of cell types, including fibroblasts, endothelial cells, smooth muscle cells, immune cells and in the chondrocytes associated with elastic cartilage and fibrocartilage (Chen et al., 2021; Strutz et al., 1995; Szabo et al., 2019; Teng et al., 2011). FSP1 performs diverse functions in the nucleus, cytoplasm, and extracellular space, acting both intracellularly and as an extracellular paracrine molecule affecting cell motility, viability, differentiation, and contractility (Ackerman et al., 2019; Boye and Mælandsmo, 2010; Per Björk et al., 2013; Schneider, Hansen, and Sheikh, 2008; Teng et al., 2011). Recent research has utilized FSP1 as a fibrochondrocyte marker to differentiate fibrocartilage, with a focus on the TMJ disc (Park et al., 2015; Su et al., 2020). However, the specific role of FSP1 in the structure of the TMJ remains unexplored.

In this study, we identify FSP1-expressing fibroblasts in the superficial layer of the TMJ as stem/progenitor cells crucial for condylar cartilage growth and joint homeostasis. This finding highlights the potential importance of FSP1 in TMJ development and maintenance, offering a novel model for studying osteoarthritis in the TMJ.

## Materials and Methods

### Animal preparation

*FSP1-Cre* mice (Bhowmick, 2004) were mated to *R26RmTmG* (referred to as *mTmG*) (Muzumdar et al., 2007), *R26RDTA* (referred to as *DTA*) (Ivanova et al., 2005), and *Ctnnb1*^*ex3(loxP)*^ (referred to as *βcatGOF*) (Harada, 1999) mice to generate transgenic mice for lineage tracing, lineage depletion and gain of function studies under the approval of the Ministry of Education, Youth and Sports of the Czech Republic (license # MSMT-24093/2021-3), supervised by the Expert Committee for Laboratory Animal Welfare of the Faculty of Medicine, Masaryk University, at the Laboratory Animal Breeding and Experimental Facility of the Faculty of Medicine, Masaryk University (facility license 310 #58013/2017-MZE-17214). Transgenic animals were maintained on a C57BL/6 background. This work was carried out under UK Home Office license and regulations in line with the regulations set out under the United Kingdom Animals (Scientific Procedures) Act 1986. The mice (n=80) were housed in individually ventilated or open cages, all with ambient temperature of 22°C, a 12 h:12 h light:dark cycle, and food and water *ad libitum*.

*Axin2-CreERT2;tdTom* mice were intraperitoneal (I.P) injected with tamoxifen (0.15 mg/g body weight) at P16 and culled at P18. For 5-Bromo-2’-deoxyuridine (BrdU) labelling, pregnant CD1 mice were I.P. injected with 20 mg/kg BrdU at E17.5 and E18.5, and the pups were culled at P2, P21, and P48.

### Tissue preparation for microcomputed tomography scanning and immunohistochemistry staining

Tissue samples were fixed in 4% paraformaldehyde and washed using phosphate-buffered saline (PBS). Skull samples were placed into a 19 mm tube containing 70% ethanol and scanned using a SCANCO MEDICAL µCT 50 scanner (70 kV, 114 uA, 20 um). The TMJ was reconstructed in 3D using Amira and MeshLab software.

Samples were decalcified in 0.5M ethylenediaminetetraacetic acid (EDTA), dehydrated, embedded in paraffin, and sectioned at 6 µm in the frontal plane using a Leica RM2245 microtome (Leica Biosystems). Samples were then mounted in parallel sequence on TruBOND380 slides (Matsunami).

### Histology staining and TUNEL (TdT-mediated dUTP Nick End Labelling) assay

Serial tissue sections of the TMJ were stained with trichrome (picrosirius red, alcian blue, and haematoxylin), haematoxylin and eosin (H&E) and safranin O staining for histological investigation (Appendix Fig. 1). A TUNEL assay was performed using the *In situ* apoptosis detection kit (TaKaRa, cat. #MK500) following the manufacturer’s instructions. A NanoZoomer Digital Slide scanner 2.0-HT (Hamamatsu Photonics) with NDP.view2 software was used to examine and capture the morphology of histology-stained TMJ slides.

Osteoarthritis levels in the TMJ were scored in safranin O or trichrome stained sections using the osteoarthritis research society international (OARSI) scale (Laverty et al., 2010).

### Immunofluorescence and RNAscope

Immunofluorescence staining was conducted for FSP1, SOX9, type II collagen (COL2A1), green fluorescent protein (GFP), red fluorescent protein (RFP), BrdU (Appendix Table 1). After deparaffinization using Neoclear and rehydration in a declining series of ethanol dilutions, paraffin sections were covered in trypsin at room temperature (RT) for 10 minutes for antigen retrieval. For collagen staining, tissue sections were enzymatically treated with chondroitinase ABC (0.1 unit/ml) and hyaluronidase (0.6 unit/ml) for 45 minutes at 37°C. The sections were incubated for 30 minutes at RT using a generic blocking buffer (1% BSA, 0.1% Triton X-100 in PBS). Sections were then treated overnight at 4°C with primary antibodies diluted in blocking buffer in a moisture chamber. The slides were washed and incubated with secondary antibodies diluted in blocking buffer for 2 hours at RT in dark. Nuclear counterstaining was performed using DAPI (Fluoroshield™ Sigma-Aldrich).

RNAscope from Advanced Cell Diagnostics (RNAscope® Multiplex Fluorescent Detection Reagents Kit v2) was used following the manufacturer’s instructions. *Axin2, Lgr5, Gli1, Scleraxis* probes were utilized (Appendix Table 1). Negative control staining was carried out (Appendix Fig. 2). Slides were imaged on a confocal microscope (ZEISS LSM 980) and ZEISS Apotome.2. Experimental data was analysed and quantified using ImageJ and Qupath-0.5.1.

### Collagen hybridizing peptide staining

Tissue sections were deparaffinized and rehydrated. 5% Goat serum in PBS was added and incubated for 30 minutes at RT to block nonspecific binding. The biotin conjugated collagen hybridizing peptide (B-CHP) solution (15 µM, 50 µL per section) in 1% BSA was heated in the oven at 80°C for 5 minutes and immediately incubated on ice for 15 seconds to quench the hot solution to RT. The B-CHP solution was quickly pipetted onto each slide within 1 minute and incubated overnight at 4°C (Hwang et al., 2017). The slides were treated with Alexa Fluor^®^ 647-streptavidin solution (0.005 mg/ml) in 1% BSA for 1 hour at RT. Nuclear counterstaining was performed using DAPI.

### Statistical analysis

All experiments were replicated at least three times. Statistical analyses were performed using GraphPad Prism 10 software. For statistical comparisons among the four groups, a one-way analysis of variance (ANOVA) followed by Tukey’s post hoc test, and a two-way ANOVA followed by Sidak’s or Tukey’s multiple comparison test were utilized. To compare the control and mutant groups, an unpaired t-test was used. Data were expressed as the mean ± standard deviation. A statistically significant difference was defined as *p-*values <0.05.

## Results

### FSP1 is progressively expressed in the superficial zone of the TMJ condyle

The expression of FSP1 was followed during murine postnatal development from birth to postweaning (Fig. 1A-E). At postnatal day 1 (P1), only a few FSP1-positive cells were observed associated with the top layers of the condyle at a distance from the SOX9 and Collagen II positive layers (Fig. 1F), with this number increasing as the condyle matured (Fig. 1G-I). Following weaning at P28, robust FSP1 expression was observed reaching the SOX9-positive chondrogenic zone in the condyle (Fig. 1J). At embryonic day (E)17.5, no FSP1 expression was observed in the condyle, highlighting FSP1 as a postnatal marker of condylar fibroblasts (Fig. 1K).

**Figure 1.**
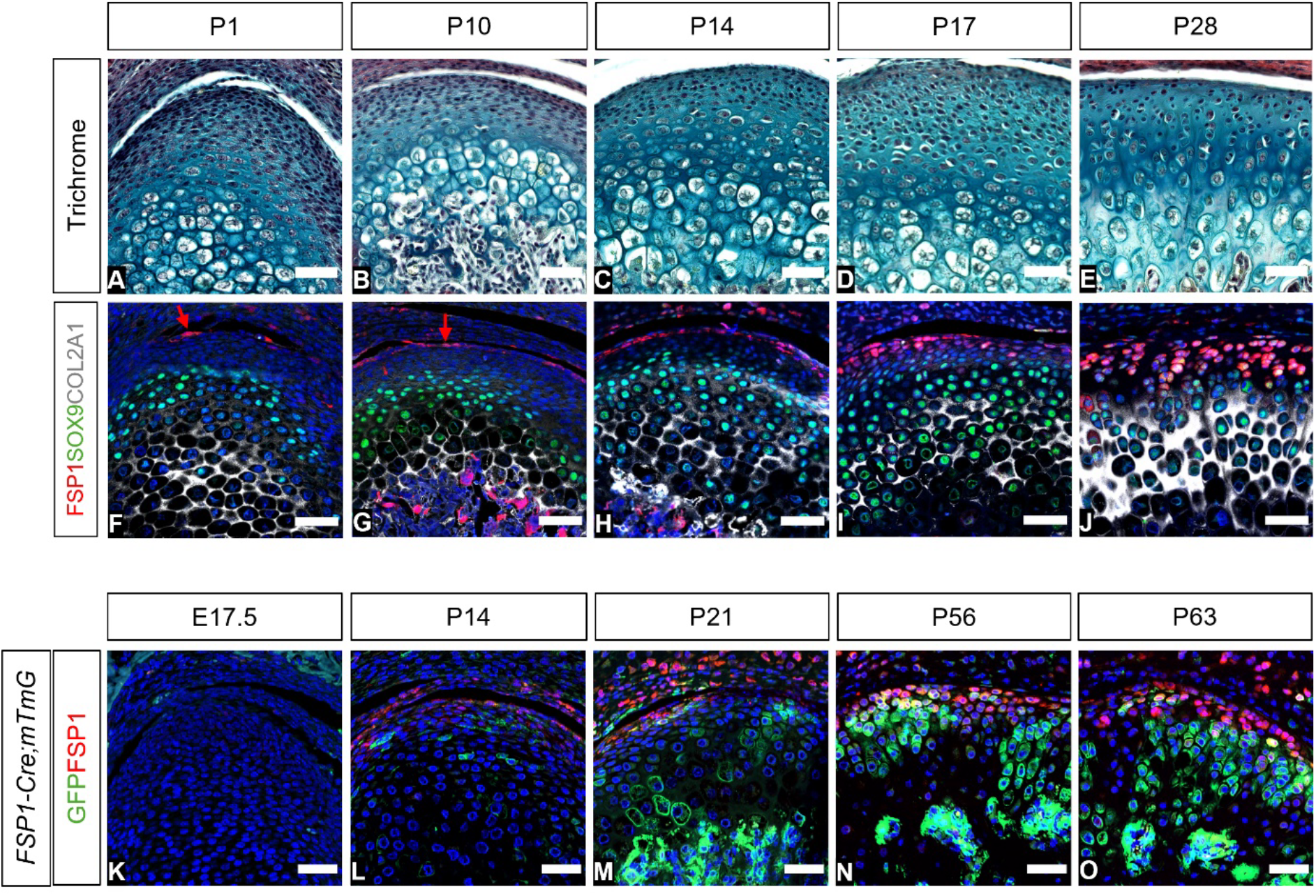
Expression and contribution of FSP1 cells in the mouse mandibular condyle across development and growth. (A-E) Picrosirius red – alcian blue trichrome and (F-J) Immunofluorescence staining for FSP1 (red), SOX9 (green), COL2A1 (grey), and DAPI (blue) illustrating the expression pattern of FSP1 during postnatal growth in CD1 mice (n=3). (F,G) Arrows indicate FSP1-positive cells (red) in the top layers of the condyle. (K-O) Lineage tracing experiment in *FSP1-Cre;mTmG* mice (n=3). Immunofluorescence staining for FSP1 was used to illustrate current FSP1 expression (red) with lineage tracing of these cells (GFP, green), and DAPI (blue). Scale bar: 50 µm. E, embryonic stage; P, postnatal stage; FSP1, fibroblast specific protein 1; GFP, green fluorescent protein.

Expression of FSP1 in the superficial zone of the condyle suggested an association with the proposed condyle stem cell niche (Embree et al., 2016). To investigate the contribution of the FSP1 positive cells to condyle growth, we employed *FSP1-Cre;mTmG* mice to lineage trace the FSP1 population. Immunostaining for FSP1 was used to compare current FSP1 expression (red) with lineage tracing of these cells (green) (Fig.1K-O). Postnatally, FSP1-positive cells were observed in the top layer of the condyle, while FSP1-lineage cells expanded into the main body of the condyle (Fig. 1L,M). By P56 and P63, the *FSP1Cre*-driven reporter expression had greatly expanded, with long clones of cells reaching the hypertrophic zones and into the ossified region (Fig. 1N,O). This expansion indicated a substantial contribution of the FSP1 lineage to the mature growth of the condyle.

### Lgr5 and Axin2 expression decrease as FSP1 switches on

Canonical Wnt signalling is crucial for maintaining fibrocartilage homeostasis (Embree et al., 2016, Ruscitto et al., 2023). *In situ* hybridization (RNAScope) and immunofluorescence was employed to investigate the relationship between *Axin2, Lgr5*, and FSP1 from E16.5 to P28. During embryogenesis, *Lgr5* was strongly expressed in the TMJ disc and partially in the condyle, with its expression decreasing postnatally (Fig. 2A-E). Similarly, *Axin2*-positive cells were abundant throughout the condyle from the embryonic stage, showing a rapid decline in the superficial layers, particularly after weaning, as FSP1 levels increased (Fig. 2A-F, Appendix Fig. 3). Some cells expressed both *Axin2* and FSP1 postnatally. This was confirmed by analysis of *Axin2-CreERT2;tdTom* mice injected with tamoxifen at P16 and collected at P18 (Fig. 2G), highlighting that FSP1 cells are in the *Axin2* lineage (Fig. 2H,I).

**Figure 2.**
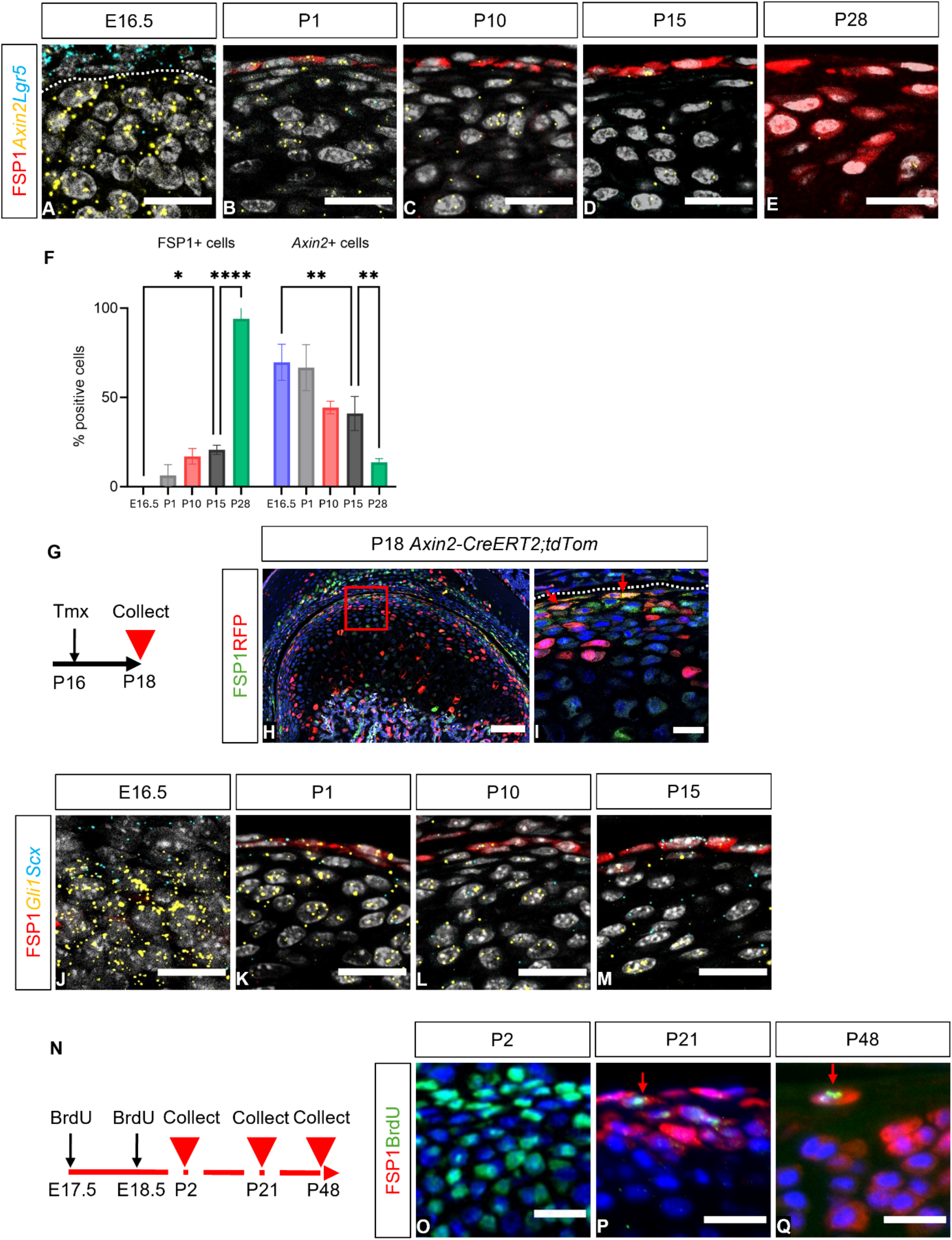
The FSP1 population co-expresses putative stem cell markers. (A-E) Dual immunofluorescence and RNAscope staining for FSP1 protein (red), *Axin2* mRNA (yellow), *Lgr5* mRNA (cyan), and DAPI (grey) between E16.5 and P28 in CD1 mice (n=3). (A) Dash line indicates condyle surface. (F) Quantification of FSP1-positive cells and *Axin2*-positive cells during condyle growth. Error bar = ±1S.D.; n=3; two-way ANOVA flowed by Tukey’s multiple comparisons test (*P<0.05, **P<0.01, ****P<0.0001). (G) Tamoxifen was administered to *Axin2-CreERT2;tdTom* mice at P16 and collected at P18 (n=3). (H) Immunofluorescence staining for FSP1 (green), RFP (red), and DAPI (blue). (I) The box shows a zoomed-in image; arrows indicate FSP1-positive cells (green) co-expressing with *Axin2* (red). (J-M) Dual immunofluorescence and RNAscope for FSP1 (red), *Gli1* mRNA (yellow), *Scx* mRNA (cyan), and DAPI (grey) in different stages of CD1 mice (n=3). (N) BrdU was administered to CD1 mice at E17.5, E18.5 (n=3). BrdU-positive label retaining cells (LRCs) were chased in P2, P21, and P48. (O-Q) Immunofluorescence staining for BrdU (green), FSP1 (red), and DAPI (blue). (P,Q) Arrows indicate FSP1-positive cells (red) co-stained with BrdU LRCs (green). Scale bar in A-E and I-Q: 20 µm, scale bar in H: 100 µm. ANOVA, analysis of variance; BrdU, 5-Bromo-2’-deoxyuridine; RFP, red fluorescent protein; Tmx, tamoxifen; Scx, scleraxis.

### A subset of FSP1 cells overlap with putative stem cell markers and are label retaining

A number of fibroblast stem cell markers have been proposed in the condyle and other fibroblast populations (Ma et al., 2021, Lei et al., 2022). Interestingly, a sub-population of FSP1-positive cells also expressed *Gli1* and *Scleraxis*, suggesting these cells had stem cell-like properties (Fig. 2J-M). Retention of a BrdU label is widely used as a hallmark of stem cells (Sottocornola and Lo Celso, 2012). To identify whether the FSP1 positive cells were label retaining, pregnant mice were injected with BrdU during TMJ morphogenesis at E17.5 and E18.5 (Fig. 2N). Confirming uptake of the label, BrdU-positive cells were observed in most condylar cells at P2 (Fig. 2O). The cells of the condyle proliferated rapidly during postnatal development, resulting in a reduce number of BrdU-positive cells at P21 and P48. Importantly, immunostaining results revealed that all BrdU-positive cells in the top layers of the condyle were FSP1-positive (Fig. 2P,Q), suggesting that FSP1 labels a novel stem/progenitor cell population in the TMJ.

### Removal of the superficial layers by Fsp1Cre-driven diphtheria toxin led to loss of condyle structure

To follow the role of the superficial layers of growth and homeostasis of the condyle, these layers were selectively ablated using the *FSP1-Cre;DTA* mouse model. In the *FSP1-Cre;DTA* mouse, *FSP1*-expressing cells were selectively exposed to diphtheria toxin and die. The toxin cannot spread to neighbouring cells as mice do not contain the receptor for this toxin (Chang and Neville, 1978). In Cre-negative *DTA* littermate control 4-week-old and 16-week-old mice, the condyle showed a mature pattern, with FSP1-positive fibroblasts reaching to the SOX9 and COL2A1 positive zones (Fig. 3A-D,3I-L). At 4 weeks, conditional ablation of FSP1-cells led to a loss of the fibrocartilage layer of the condyle (Fig. 3E,F) with positive TUNEL cells observed in this area (Fig. 3G). A reduction in the number of FSP1- and SOX9-positive cells was also noted by immunofluorescence (Fig. 3H). At 16 weeks, a clear reduction in FSP1 expression was observed in mutants compared to littermates, confirming the loss of a large proportion of the FSP1 population (Fig. 3P). As predicted, the *FSP1-Cre;DTA* mice exhibited nearly complete loss of condylar cartilage. H&E staining highlighted the pronounced transformation in the layers of the condyle, culminating in a severe osteoarthritic phenotype (Fig. 3M,N). The condylar surface became disrupted and uneven. The disruption of the superficial layers triggered ectopic cartilage formation, resulting in elevated production of SOX9 along the surface of the condyle (Fig. 3P). Following FSP1 cell ablation, we observed disruptions in ECM pattern, characterized by diminished and disorganized distribution of collagen II (Fig. 3P), and increased mean intensity of B-CHP within the superficial layer of the condyle (Fig. 3Q,S,U, Appendix Fig. 4). These findings indicate heightened collagen remodelling activity. Scoring of the phenotype using the OARSI scale, showed grade 6 osteoarthritis in all *FSP1-Cre;DTA* mice at 16-weeks (Fig. 3V). Loss of the most superficial layers of the condyle, therefore, led to osteoarthritic remodelling of the TMJ.

**Figure 3.**
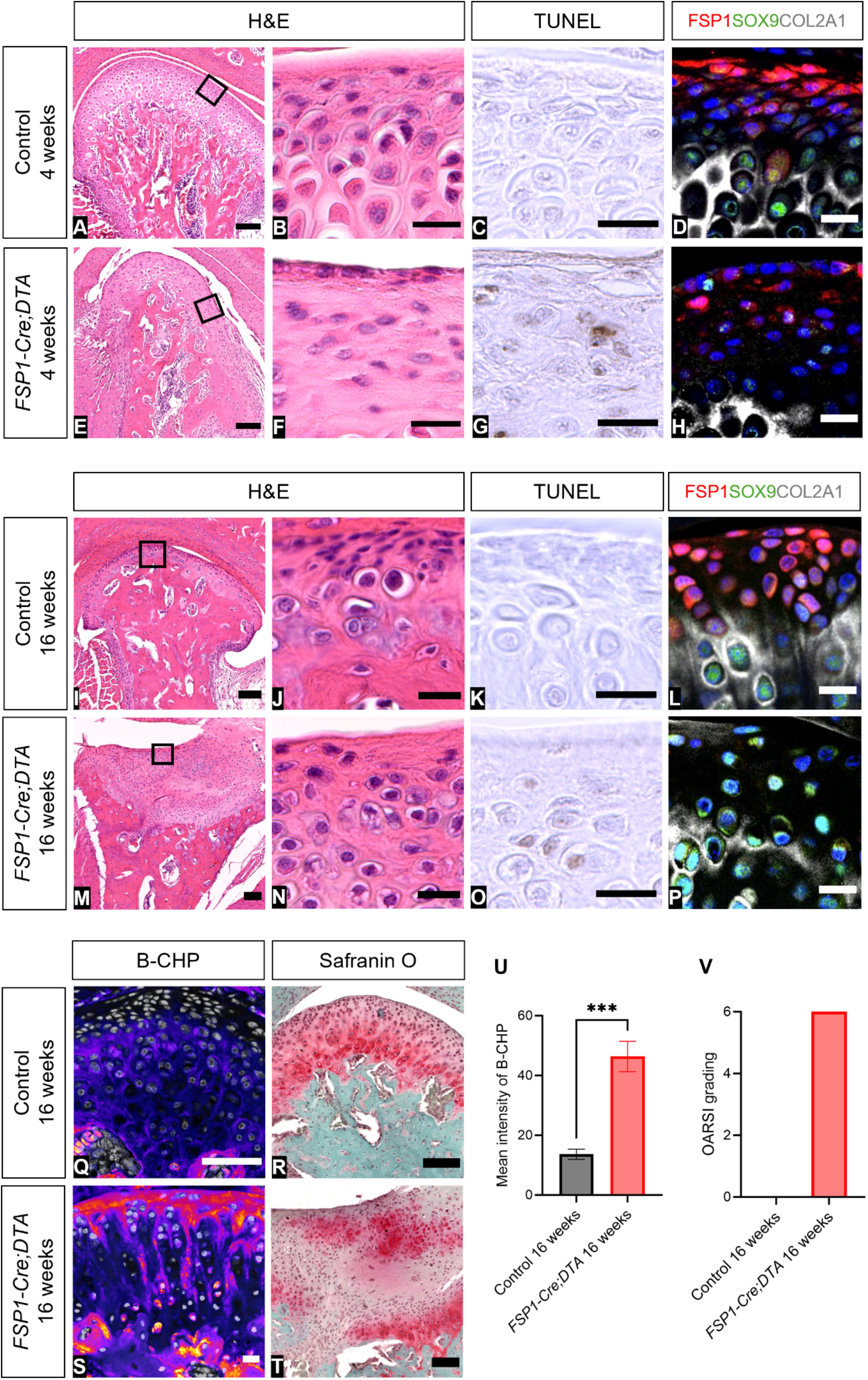
Removal of the superficial layers by *FSP1-Cre-*driven DTA led to loss of condylar structure. (A-H) and (I-P) H&E staining, TUNEL assay, and immunofluorescence staining in 4-week-old (n=3) and 16-week-old (n=6) *FSP1-Cre;DTA* mice, respectively, along with the Cre-negative *DTA* littermate controls. (A,E,I,M) H&E staining shows condyle morphology. (B,F,J,N) The box shows a zoomed-in image of superficial layer of the condyle. (C,G,K,O) TUNEL-positive cells are indicated by a brown colour cell. (D,H,L,P) Immunofluorescence staining for FSP1 (red), SOX9 (green), COL2A1 (grey), and DAPI (blue). (Q,S) 16-week-old *FSP1-Cre;DTA* mice and Cre-negative *DTA* littermate controls were stained with immunofluorescence staining for B-CHP (fire) and DAPI (grey), and (R,T) Safranin O staining. (U) Mean intensity of B-CHP was analysed using ImageJ software. Error bar = ±S.D.; n=3 unpaired t test (***P<0.001). (V) OARSI grading was blindly scored by observers (n=3). Scale bar B-D, F-H, J-L, N-P, Q-S: 20 µm, scale bar A,E,I,M,R,T: 100 µm. B-CHP, biotin conjugated collagen hybridizing peptide; H&E, haematoxylin and eosin; OARSI, osteoarthritis research society international.

### Loss of FSP1-cells impacted TMJ shape and growth of the dentary

Histological examination highlighted a significant alteration in the morphology of the condyle after ablation of the FSP1-expressing cells. To assess the impact on the condylar process and dentary bone, we employed µCT scanning. 4-week-old and 16-week-old littermate controls exhibited a normal TMJ structure, displaying a wide anterior and narrow posterior aspect (Fig. 4A,B). In contrast, in 4-week-old *FSP1-Cre;DTA* mice, a significant enlargement of the condylar head was observed in the posterior orientation (Fig. 4C). By 16 weeks, the posterior condyle was further enlarged, with the condyle having a concave surface associated with ectopic mineralised tissue (Fig. 4D). The growth of the dentary was impacted by loss of the FSP1 cells, with significant decreases in length from the condyle to the lingula and between the condyle and angular process (Fig. 4E,F). Loss of the superficial layer thus impacted the normal growth of the dentary.

**Figure 4.**
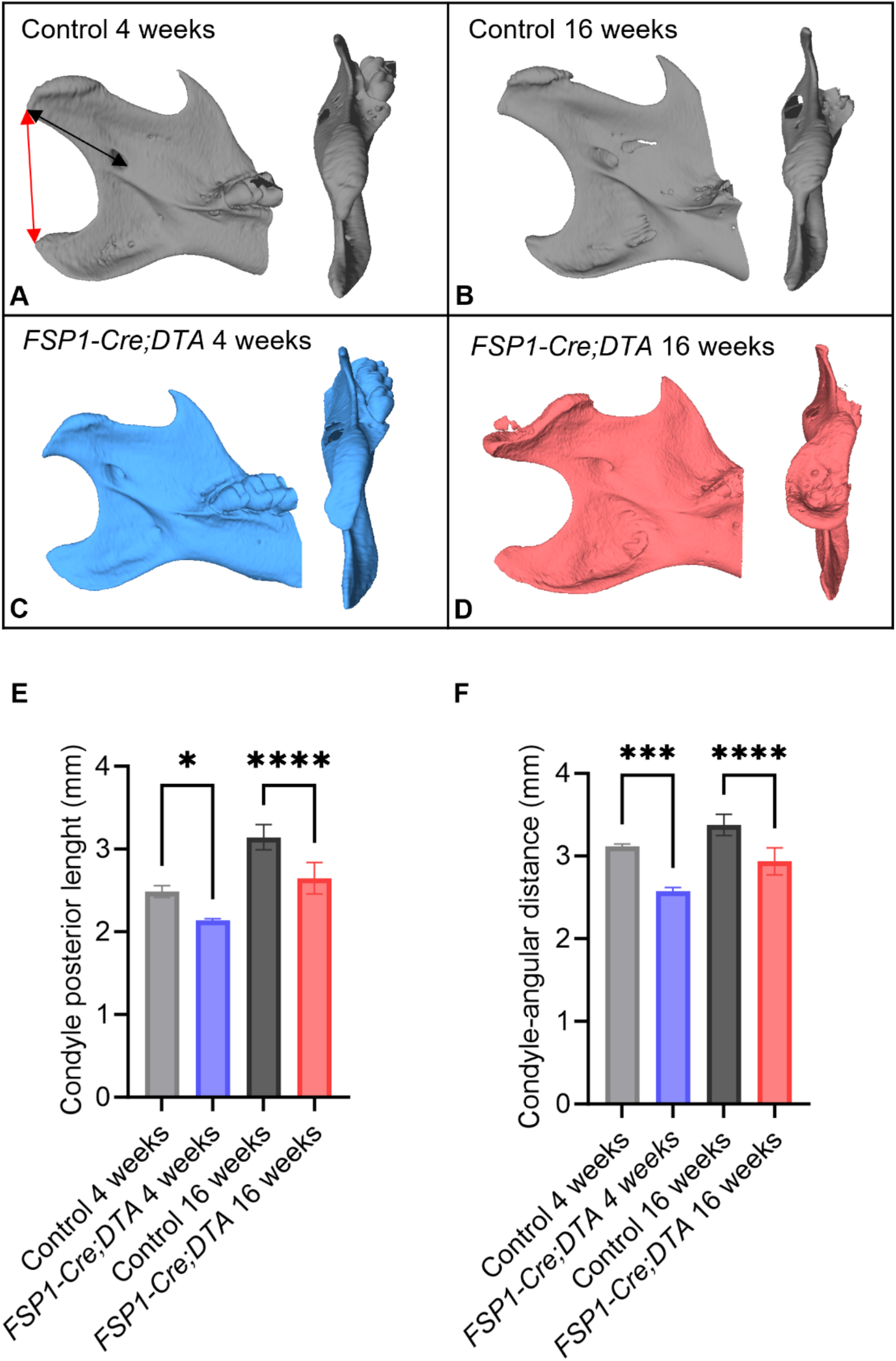
Loss of *FSP1*-cells impacted TMJ shape and growth of the dentary. (A-D) 3D reconstructions µCT scans of the condylar proximal mandible in superior view and internal-lateral view in 4-week-old (blue, n=3) and 16-week-old (pink, n=6) *FSP1-Cre;DTA* mice, respectively, along with the Cre-negative *DTA* littermate controls (grey). (A)The black arrow indicates the condyle posterior length, which is the distance from the most caudal point of the condylar process to the most rostral and ventral point of the mandibular foramen (anterior point of lingula). (A) The red arrow indicates the condyle-angular distance, which is the distance from the most caudal point of the condylar process to the tip of the angular process of the mandible. (E,F) Measurements of the condyle posterior length and the condyle-angular distance using Amira software. Error bar = ±S.D.; one-way ANOVA followed by Tukey’s multiple comparisons test (*P<0.05, ***P< 0.001, ****P < 0.0001).

### FSP1 expression was negatively regulated by canonical Wnt activity, resulting in loss of hyaline cartilage

Loss of *Axin2*, concurrent with upregulation of FSP1 expression (Fig. 2F), suggested *Axin2* expression might inhibit FSP1. To test this, we utilized *FSP1-Cre* driven β-catenin gain of function mice to constituently activate Wnt/β-catenin signalling in FSP1-expressing cells. 12-week-old *FSP1-Cre;βcatGOF* mice displayed a complete loss of condylar hyaline cartilage, evidenced by severe osteoarthritic changes when compared to Cre-negative *βcatGOF* littermates (Fig. 5A,B,D,E). The condylar surface was disrupted, with loss of FSP1- and SOX9-positive cells in the superficial zone (Fig. 5C,F). To confirm *β-catenin* function in *FSP1-Cre;βcatGOF* mice, RNAscope was used to quantify FSP1-positive and *Axin2*-positive cells, revealing an increase in *Axin2* expression associated with downregulation of FSP1 expression (Fig. 5G-I). Loss of canonical Wnt signalling in the condyle postnatally is, therefore, essential to allow upregulation of FSP1 and maintenance of the superficial layers of the condyle.

**Figure 5.**
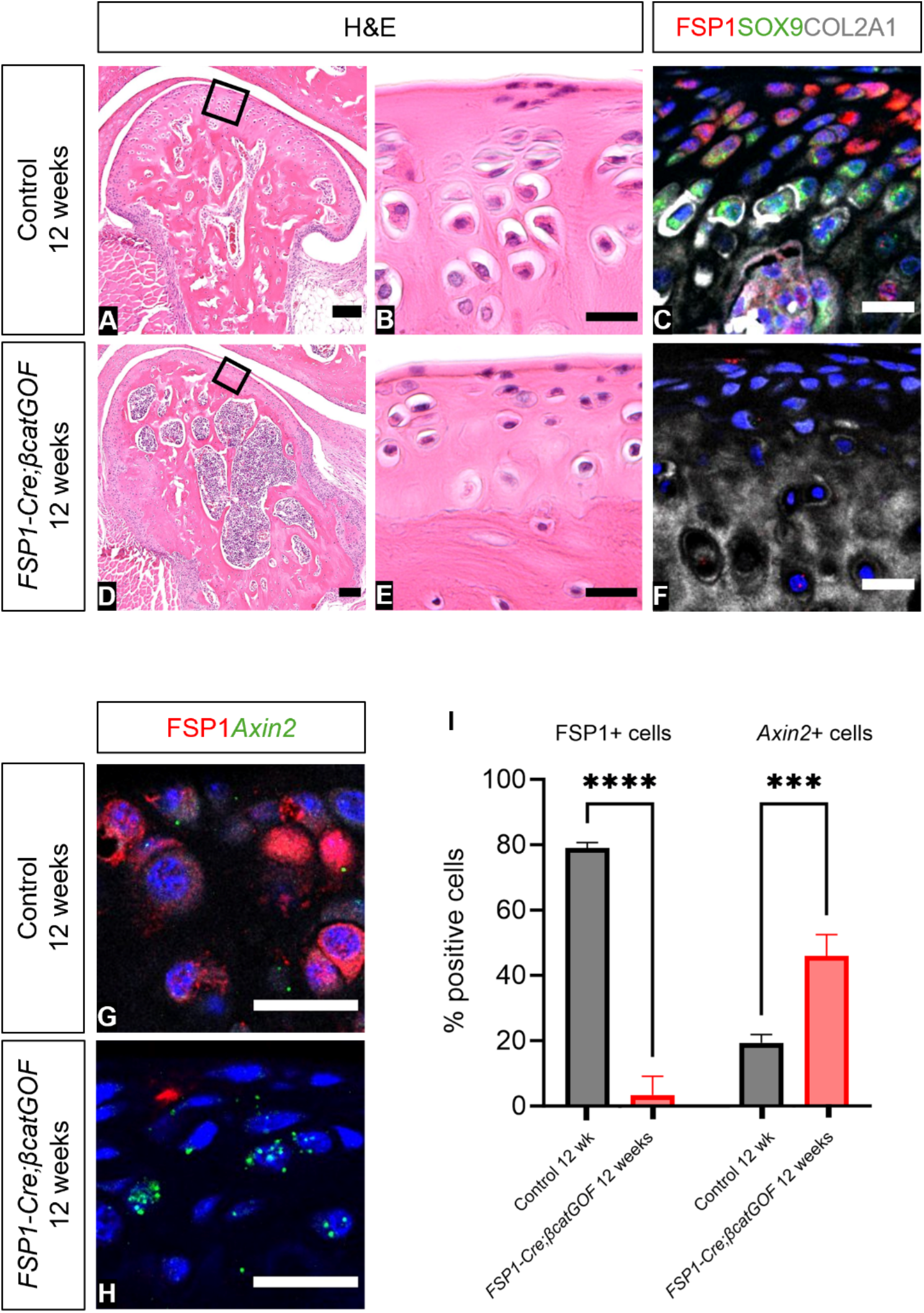
FSP1 expression is negatively regulated by canonical Wnt activity, resulting in loss of TMJ hyaline cartilage. (A-F) H&E staining and immunofluorescence staining in 12-week-old *FSP1-Cre;βcatGOF* (n=3) and Cre-negative *βcatGOF* littermate controls. (A,D) H&E staining shows condyle morphology. (B,E) The box shows a zoomed-in image of superficial layer of the condyle. (C,F) Immunofluorescence staining for FSP1 (red), SOX9 (green), COL2A1 (grey), and DAPI (blue). (G,H) Representative images of immunofluorescence with RNAscope staining for FSP1 (red), *Axin2* (green), and DAPI (blue) in 12-week-old *FSP1-Cre;βcatGOF* and Cre-negative *βcatGOF* littermate controls. (I) FSP1-positive cells and *Axin2*-positive cells were quantified using Qupath software. Error bar = ±S.D.; n=3; two-way ANOVA followed by Sidak multiple comparisons test (***P<0.001, ****P<0.0001). Scale bar B-C, E-F, G-H: 20 µm, scale bar A and D: 100 µm. *βcat*, β-catenin; GOF, gain of function.

## Discussion

We propose FSP1 as a novel marker for the superficial fibroblast layers of the mature condyle. FSP1 was induced postnatally, starting in the most superficial layer and later extending to the SOX9 chondrogenic zone of the condyle. Numbers of FSP1-expressing cells increased during development with a sharp rise and expansion of the domain around tooth eruption (Lungova et al., 2011), suggesting mechanical force may act as a trigger for upregulation. The FSP1-positive cells could be considered a stem/progenitor population for the postnatal mouse TMJ, with lineage tracing showing that they gave rise to all layers of the condylar process during postnatal development. Additionally, a subset of FSP1-expressing cells overlapped with other proposed fibroblast stem cell markers, Gli1 and Scleraxis, and were derived from Wnt-responsive *Axin2* cells. A subset of FSP1 cells were also label-retaining, indicating that the FSP1 cells marked both stem and progenitor populations. As cells exited the superficial zone and started to differentiate and turn on cartilage markers, expression of FSP1 was downregulated.

Loss of the superficial zone, by deletion of FSP1 cells, led to severe osteoarthritis of the TMJ. Loss of FSP1 cells in our model would impact both the stem/progenitor cells of the condyle and the superficial synovial cells. Our phenotype was similar to that reported in the *Pgr4* mutant mice, where lubricin production is prevented, resulting in defects in the synovial fluid (Bechtold et al., 2016). In these mutants, the presence of SOX9-positive cells, coupled with abnormalities in cellular integrity and morphology, led to the formation of osteophytes over time (Bechtold et al., 2016). Loss of the FSP1 cells led to upregulation of SOX9, extensive remodelling of the ECM, and ectopic cartilage formation. Chondrocytes play a pivotal role in maintaining cartilage matrix homeostasis, and any compromise in their activity and survival can disrupt this delicate balance, hastening the progression of osteoarthritis (Lu et al., 2022; Embree et al., 2010). The OA phenotype was consistent with previous studies that have demonstrated that degradation of collagen II can promote chondrocyte hypertrophy, thereby accelerating the development of TMJ osteoarthritis (Lian et al., 2019). Interestingly, recent research has shown that FSP1 plays a role in producing collagen I postnatally, a critical factor for bone development, maintenance, and repair (Chen et al., 2021). When collagen I was deleted in FSP1-positive cells, it led to osteogenic imperfecta-like symptoms in adult mice, characterized by spontaneous fractures and impaired bone healing (Chen et al., 2021).

Expression of FSP1 increased postnatally at the same time as active Wnt signalling was downregulated in the superficial zone. Sustained Wnt signalling in this layer led to loss of FSP1, highlighting the importance of reducing Wnt signalling in this layer for maintenance of the stem/progenitor population. Previous studies using conditional activation of β-catenin mice have demonstrated TMJ osteoarthritis-like phenotypes, such as cartilage degradation, upregulation of collagen X, decreased cell proliferation, and increased cell apoptosis (Hui et al., 2018). Excessive Wnt signalling has also been shown to disrupt fibrocartilage homeostasis, causing degenerative changes, that lead to the deterioration of the fibrocartilage stem cell population (Embree et al., 2016). Similarly, we have shown that constitutive Wnt/β-catenin signalling specifically in the FSP1 population led to an osteoarthritic phenotype, providing a mechanism for the OA phenotype previously observed. In summary, our research highlights the importance of the FSP1-expressing superficial zone, the interplay between FSP1 and Wnt signalling, and provides valuable models for investigating TMJ osteoarthritis.

## Author contribution

AST, NA conceptualized the work, AST, NA, TT designed experiments. TT, NA, WW, DB performed experiments and analysed data. DB, NA, WW performed tissue collection. TT wrote original manuscript. NA, AST, DB, ZSK edited the manuscript. AST, NA, and ZSK acquired funding, supervised the work. All authors gave their final approval and agreed to be accountable for all aspects of the work.

