## Appendix for "FSP1 stem/progenitor cells are essential for TMJ growth and homeostasis"

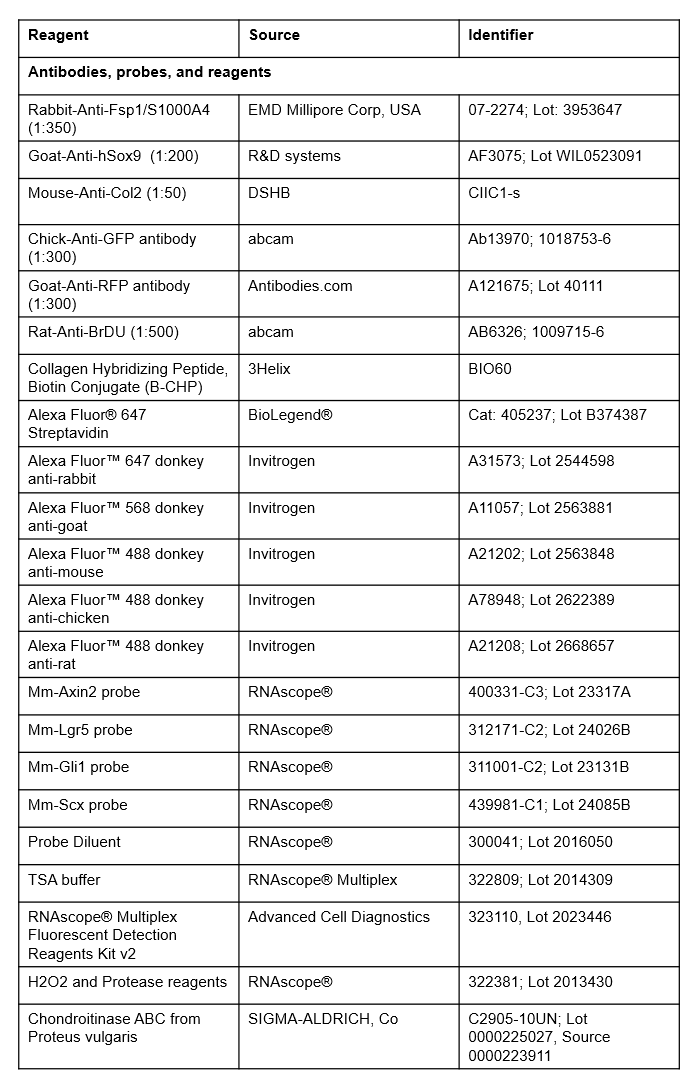
**Appendix Table1** The table shows the antibodies, probes, and reagents used for all experiments.
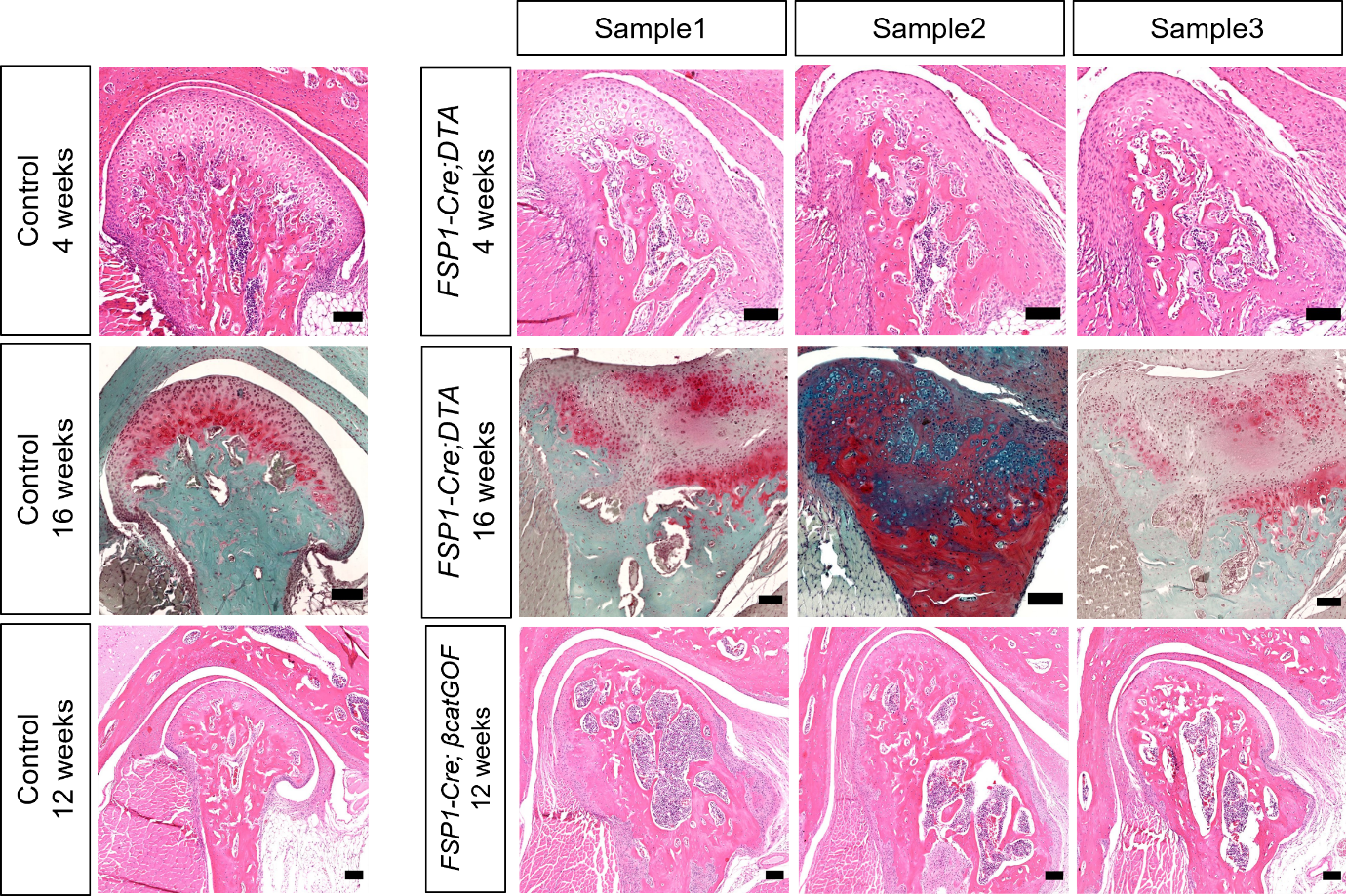
 **Appendix Figure1** Histology staining of 4-week-old *FSP1-Cre;DTA* mice, 16-week-old *FSP1-Cre;DTA* mice, and 12-week-old *FSP1-Cre;βcatGOF* mice (n=3). scale bar: 100 µm.

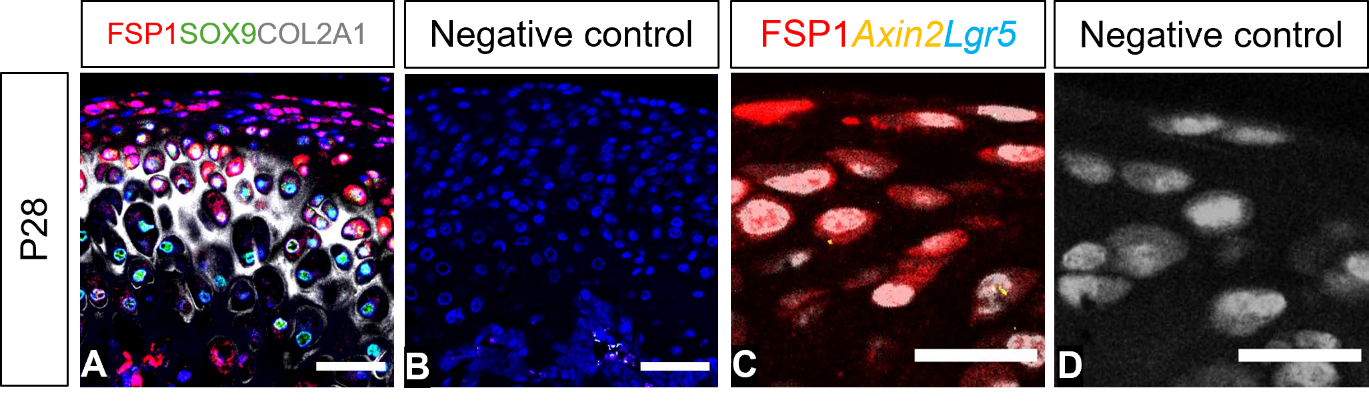

**Appendix Figure2** (A) Immunofluorescence staining for FSP1 (red), SOX9 (green), COL2A1 (grey), and DAPI (blue) in P28. (B) a negative control slide of immunofluorescence staining. (C) Dual immunofluorescence and RNAscope staining for FSP1 protein (red), *Axin2* mRNA (yellow), *Lgr5* mRNA (cyan), and DAPI (grey) in P28. (D) a negative control slide of dual immunofluorescence and RNAscope staining. Scale bar A-B: 50 µm, scale bar in C-D: 20 µm.

**Appendix Figure3** Quantification of FSP1-positive cells and *Axin2*-positive cells during condyle growth. Error bar = ±S.D.; n=3; two-way ANOVA flowed by Tukey’s multiple comparisons test (*P<0.05, **P<0.01, ***P<0.001, ****P<0.0001).
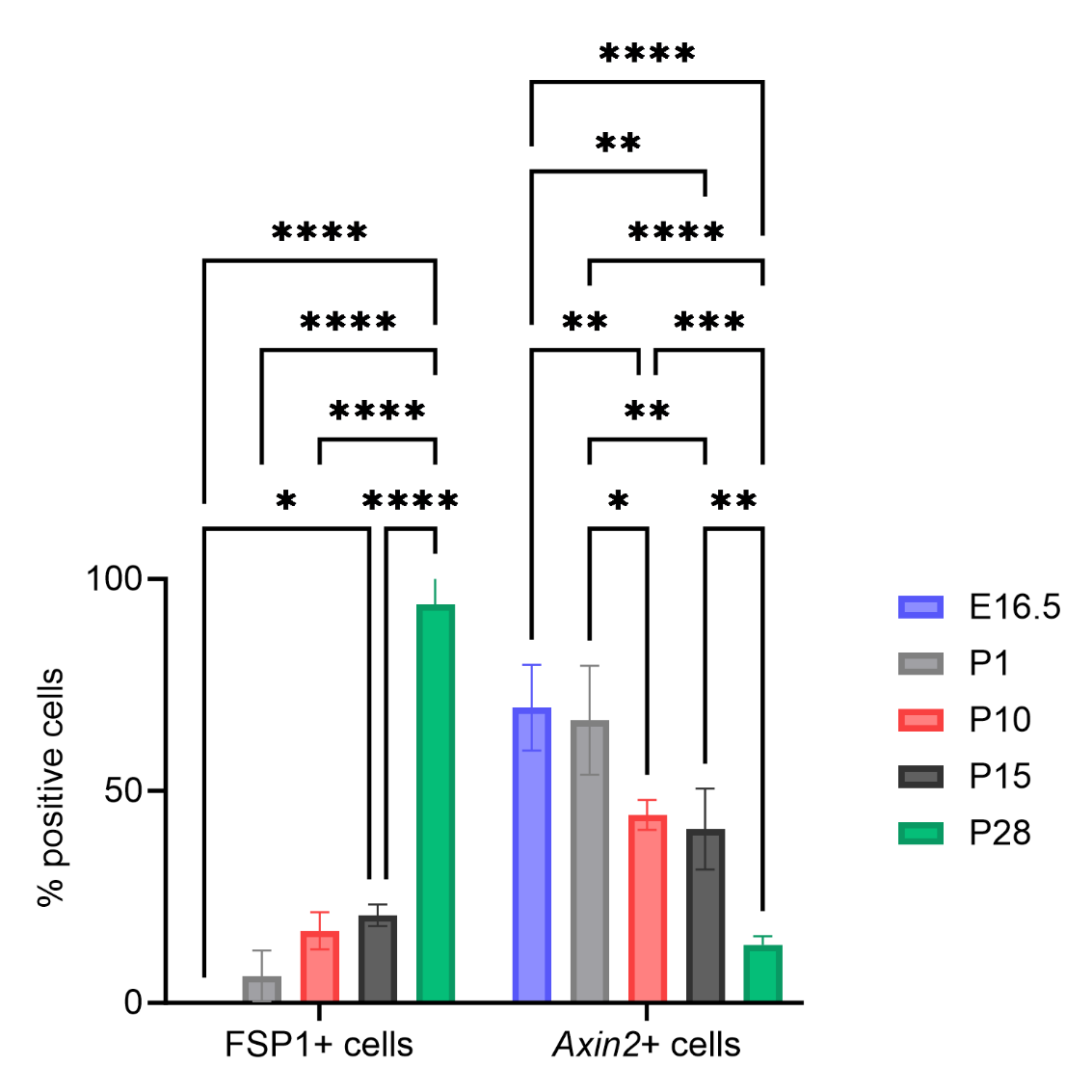

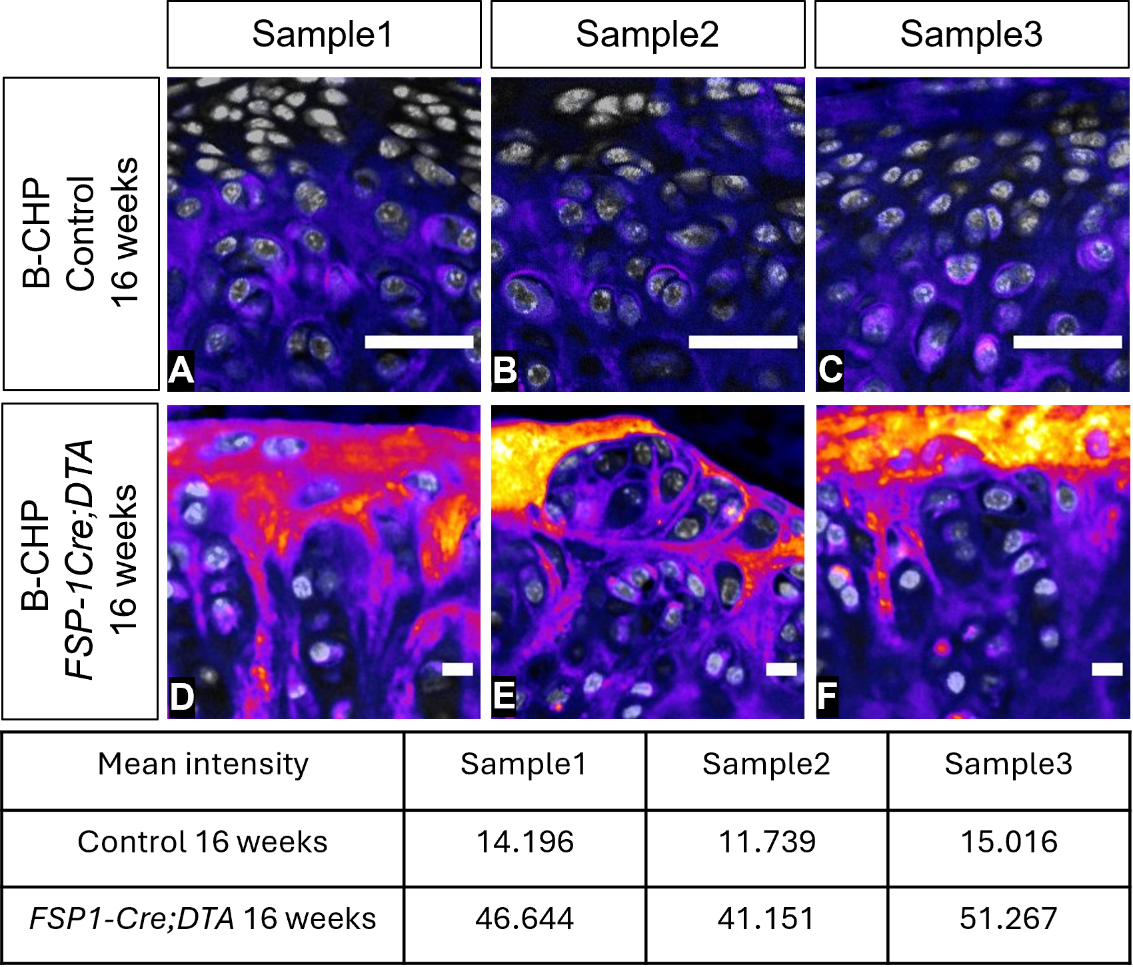

**Appendix Figure4** (A-F) 16-week-old *FSP1-Cre;DTA* mice and the Cre-negative *DTA* littermate controls were stained with immunofluorescence staining for B-CHP (fire) and DAPI (grey). The table below shows the mean intensity of B-CHP. scale bar in A-F: 20 µm.
